# Prediction pipeline for discovery of regulatory motifs associated with *Brugia malayi* molting

**DOI:** 10.1101/781930

**Authors:** Alexandra Grote, Yichao Li, Canhui Liu, Denis Voronin, Adam Geber, Sara Lustigman, Thomas R. Unnasch, Lonnie Welch, Elodie Ghedin

**Affiliations:** Department of Biology, Center for Genomics and Systems Biology, New York University, New York, NY 10003, USA; School of Computer Science and Electrical Engineering, Ohio University, Athens OH 45701, USA; Center for Global Infectious Disease Research, University of South Florida, Tampa, FL 33612, USA; Laboratory of Molecular Parasitology, Lindsley F. Kimball Research Institute, New York Blood Center, New York, NY 10065, USA; Department of Epidemiology, College of Global Public Health, New York University, New York, NY 10003, USA

## Abstract

Filarial nematodes can cause debilitating diseases in humans. They have complicated life cycles involving an insect vector and mammalian hosts, and they go through a number of developmental molts. While whole genome sequences of parasitic worms are now available, very little is known about transcription factor (TF) binding sites and their cognate transcription factors that play a role in regulating development. To address this gap, we developed a novel motif prediction pipeline, Emotif Alpha, that integrates ten different motif discovery algorithms, multiple statistical tests, and a comparative analysis of conserved elements between the filarial worms *Brugia malayi* and O*nchocerca volvulus*, and the free-living nematode *Caenorhabditis elegans.* We identified stage-specific TF binding motifs in *B. malayi*, with a particular focus on those potentially involved in the L3-L4 molt, a stage important for the establishment of infection in the mammalian host. Using an *in vitro* molting system, we tested and validated three of these motifs demonstrating the accuracy of the motif prediction pipeline.

## Introduction

*Brugia malayi* is a mosquito-borne filarial nematode and one of the causative agents of lymphatic filariasis, commonly known as elephantiasis. Currently, 856 million people in 52 countries require preventative chemotherapy to stop the spread of the disease (Gordon et al. 2018). Transmission occurs when the mosquito vector introduces infective third-stage larvae (L3) during their blood meal. The larvae then migrate to the lymphatic vessels where they molt twice and develop into adults. Over their lifespan adult females produce millions of microfilariae (immature larvae) that circulate in the blood, allowing for continued transmission. Chronic lymphatic filariasis can cause permanent and disfiguring damage, characterized by lymphoedema (tissue swelling) and elephantiasis (tissue thickening) of the lower limbs, and hydrocele (scrotal swelling).

Over the past decade, a few parasitic nematode genomes have been sequenced, including *B. malayi* (Ghedin et al., 2007), *Loa loa* (Desjardins et al. 2013), and *Onchocerca volvulus* (Cotton et al. 2016). Transcriptomic experiments have helped quantify differentially expressed genes and their biological implications (Bennuru et al., 2016; Choi et al., 2011; Grote et al, 2017; Kariuki, Hearne, & Beerntsen, 2010; Li, Wang, Rush, Mitreva, & Weil, 2012). However, little is known about how these genes are regulated through cis-regulatory motifs. Motifs that have been characterized in *B. malayi* and that are available in the CIS-BP database (Weirauch et al. 2014) are purely bioinformatic predictions based on transcription factor binding site (TFBS) homology. *De novo* DNA motif discovery is an effective bioinformatic method for studying transcriptional gene regulation (Dieterich & Sommer, 2008), and a number of motif discovery methods and tools currently exist. These include expectation-maximization methods, such as MEME (Bailey et al. 2006) and Improbizer (Ao et al. 2004); Gibbs sampling methods, such as BioProspector (Liu et al. 2001) and MotifSampler (Thijs et al. 2002); k-mer enumeration methods such as Weeder (Pavesi, et al. 2004), DME (Smith et al. 2005), and DECOD (Huggins et al. 2011); ensemble methods such as W-ChIPMotifs (Jin et al. 2009), and GimmeMotifs (Heeringen & Veenstra, 2011); and deep learning methods such as DanQ (Quang & Xie, 2016) and DeepFinder (Lee et al. 2018). Based on the input types, motif discovery approaches can also be classified as either generative or discriminative. Generative motif discovery models use pre-defined background models (e.g., the Hidden Markov Model), while discriminative motif discovery models need to explicitly specify a set of background sequences. In this study, we developed Emotif Alpha that integrates a number of the current methods based on the aforementioned models and filters the motifs using a Z-test, random forest feature importance, and sequence homology.

Gene promoter regions play a crucial role in gene regulation yet remain largely uncharacterized in *B. malayi*. Among the very few promoters that have been previously described and validated in *B. malayi* is that of Heat Shock Protein 70 (HSP70) (Shu et al. 2003). A previous study showed that while the regulatory domains of the HSP70 promoter were similar to other eukaryotes, the core promoter domains appeared to be distinct (Higazi et al., 2005). And nothing is known about motifs regulating developmentally expressed genes in *B. malayi*. There is thus a need for systematic identification, annotation, and experimental validation of *B. malayi* promoter motifs associated with gene regulation to better characterize filaria gene expression patterns. To better understand how promoter elements regulate stage-specific gene expression, we performed differential gene expression analysis of the L3 to L4 molt, the first developmental step important for the establishment of infection in the mammalian host, and motif discovery using the Emotif Alpha pipeline. Several promoter motifs appeared to be associated with the regulation of the L3 to L4 molt. Our results provide an initial overview of the putative regulatory mechanisms in the filariae that could be targeted using novel intervention strategies for control.

## Results

### Stage-specific expression of serpins, peptidases, cysteine protease inhibitors, and structural constituents of the cuticle during the L3 to L4 molt

Since the L3 to L4 molt is of particular interest because it corresponds to the life cycle stage when infective larvae establish infection, we focused in this study on identifying genes that are differentially expressed during this unique process. We used RNA-seq to profile transcription at different time points during the molt, collecting samples from the infective L3 (from mosquitoes), L3 Day 6, and L3 Day 9 worms recovered from infected gerbils (NCBI PRJNA557263). We combined this transcriptome data with previously published L4 data (Grote et al. 2017) that corresponds to Day 14 post infection of gerbils (**Table 1**). In total, 2.36 billion reads were generated, with 1.38 billion reads mapping to the *B. malayi* genome. Each biological replicate received an average of 272 million reads, with an average of 173 million reads that were successfully mapped (**Table 1**). Of the 11,841 *B. malayi* gene models, 87.6% were expressed in at least one stage of the L3 to L4 molt (**Fig. 1**). The molting expression data shows unique stage-specific profiles for each stage of the molt with significant differential expression between days.

**Table 1:** RNA-Seq summary. The table shows the total reads sequenced and mapped in each biological replicate at each developmental stage, L3 to L4; lower case a, b, and c refer to separate biological replicates.

| Library | Total reads (millions) | Stage Total (millions) | Total Mapped Reads (millions) | Stage Total Mapped Reads (millions) | % *B. malayi* Genes Expressed |
| --- | --- | --- | --- | --- | --- |
| L3a | 294 | 763 | 266 | 639 | 84 |
| L3b | 223 |  | 204 |  |  |
| L3c | 246 |  | 169 |  |  |
| L3D6a | 508 | 971 | 131 | 477 | 82.3 |
| L3D6b | 243 |  | 152 |  |  |
| L3D6c | 220 |  | 194 |  |  |
| L3D9a | 300 | 440 | 170 | 266 | 81.2 |
| L3D9b | 140 |  | 96 |  |  |
| L4a | 104 | 195 | 47 | 96 | 72.4 |
| L4b | 90 |  | 49 |  |  |

**Figure 1:**
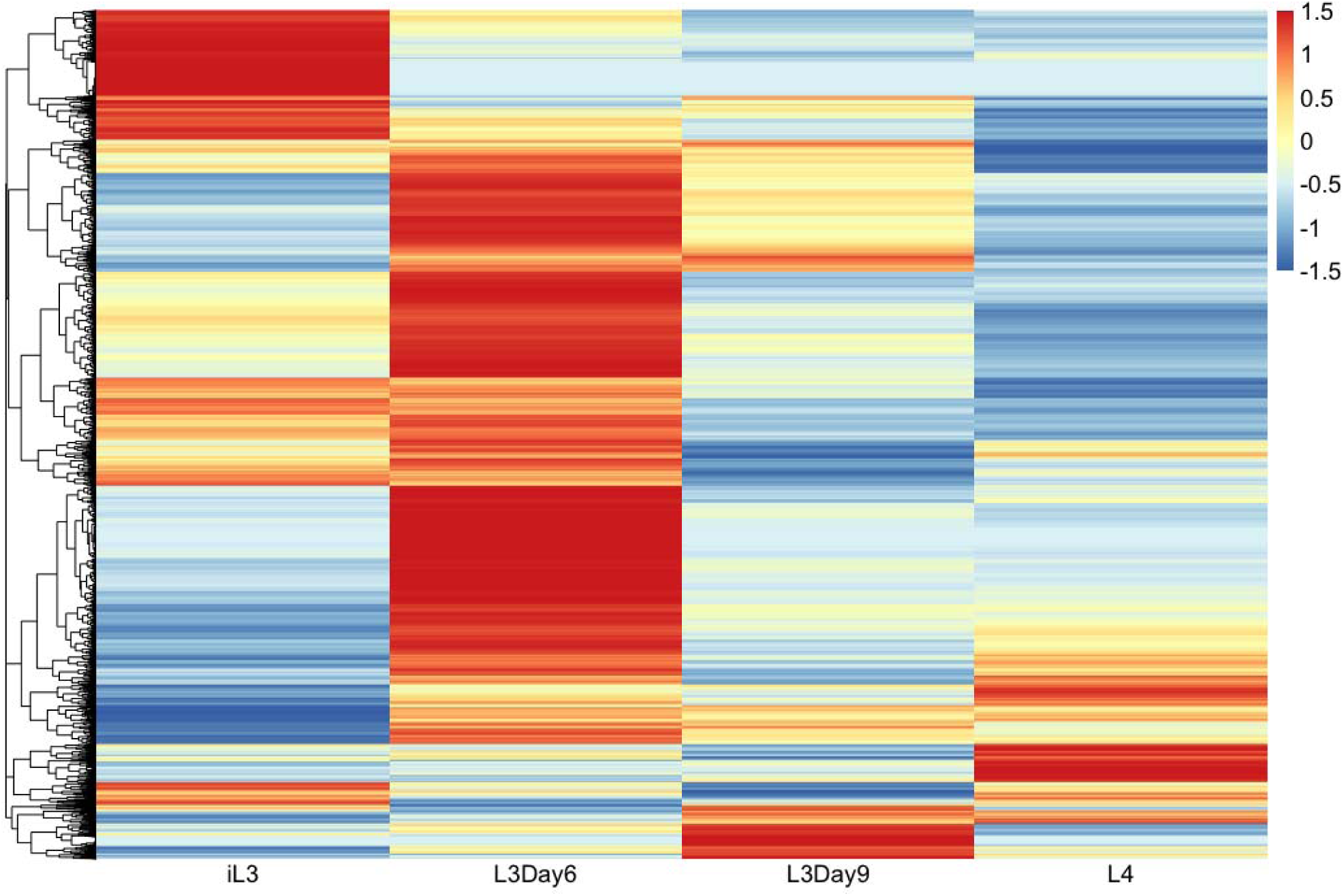
Expression of *Brugia malayi* genes during the L3 to L4 molt. Expression is in FPKMs and is Z-scale normalized by row prior to clustering. High expression is indicated by red and low expression by blue. Time-points included infective L3 larvae (iL3), L3 larvae at Day 6 of molting (L3D6), L3 larvae at Day 9 of molting (L3D9), and L4 larvae. Biological replicates have been combined.

We determined differentially expressed genes during the L3 to L4 molt using both DESeq (Anders & Huber, 2010) and EdgeR (Robinson et al. 2010) to perform pairwise comparisons between the four samples. To get a high-confidence list of differentially expressed genes, we used the consensus of the two algorithms. For the purposes of this study, we focused on the genes that were up-regulated at each stage of molting, as compared to the other stages, and did a gene-annotation enrichment analysis for each stage. We found that up-regulated genes in iL3 larvae, as compared to L3D6, L3D9, and L4, were enriched for annotations involving cysteine-type peptidase activity as well as serpin domains and serpin family proteins. Cysteine-type peptidases are essential for molting in *B. malayi* (Guiliano D.B. et al. 2004, Lustigman S. et al. 2004) and serpins are serine protease inhibitors that have previously been shown to be involved in immunomodulation and host immune evasion during infection (Zang X, et al. 2001). We identified five different cysteine-type peptidases and two cysteine-type endopeptidase inhibitors that were upregulated in iL3 larvae. By day 6 of molting, structural constituents of the cuticle, including collagen (the main component of the cuticle) were enriched in the up-regulated gene sets. We also see the up-regulation of several metalloproteases. At day 9, genes involved in signaling were enriched among the up-regulated genes, as were several different metalloproteases. At day 14 (L4 larvae), we again see an enrichment of structural constituents of the cuticle. Similarly to those enriched in L3 day 6 larvae, they are all mostly orthologs of *C. elegans* col (COLlagen) genes, which are themselves orthologs of human MARCO genes (macrophage receptor with collagenous structure). The set of structural constituents enriched at day 14 is, however, a completely unique set of collagen genes as compared to the genes observed at day 6. These stage-specific enrichments reflect the order of peptidases and structural constituents necessary for the building of a new L4 cuticle, the separation of the old L3 cuticle from the developing L4 cuticle, and the shedding of the old L3 cuticle.

### Identification of 12 motifs associated with transcription factor binding that are enriched in the L3 to L4 molt

To better understand the regulatory program of *B. malayi* during the L3 to L4 molt, we analyzed statistically over-represented DNA motifs in regions upstream of genes that were upregulated during molting. To do so, we developed a motif identification pipeline called Emotif Alpha (**Fig. 2**). First, we used the transcriptome data from the different stages of the L3 to L4 molt to generate lists of genes up-regulated at each stage of the molt using pair-wise comparisons. We then did a motif discovery analysis on each gene set using a combination of three motif discovery tools: GimmeMotifs, DME, and DECOD. GimmeMotifs is an ensemble of generative motif discovery tools—including Homer (Heinz et al. 2010), AMD (Shi et al. 2011), BioProspecter, MDmodule (Conlon et al. 2003), MEME, Weeder, GADEM (Li et al. 2009), and Improbizer—while DME and DECOD are discriminative motif discovery tools. We did a discriminative motif discovery analysis by randomly selecting background promoter region sets from all *B. malayi* genes, excluding the differentially expressed genes. These background sets are three times larger than the foreground sets. We selected motif lengths between 6- and 15-mer. In total, we identified 20,025 motifs.

**Figure 2:**
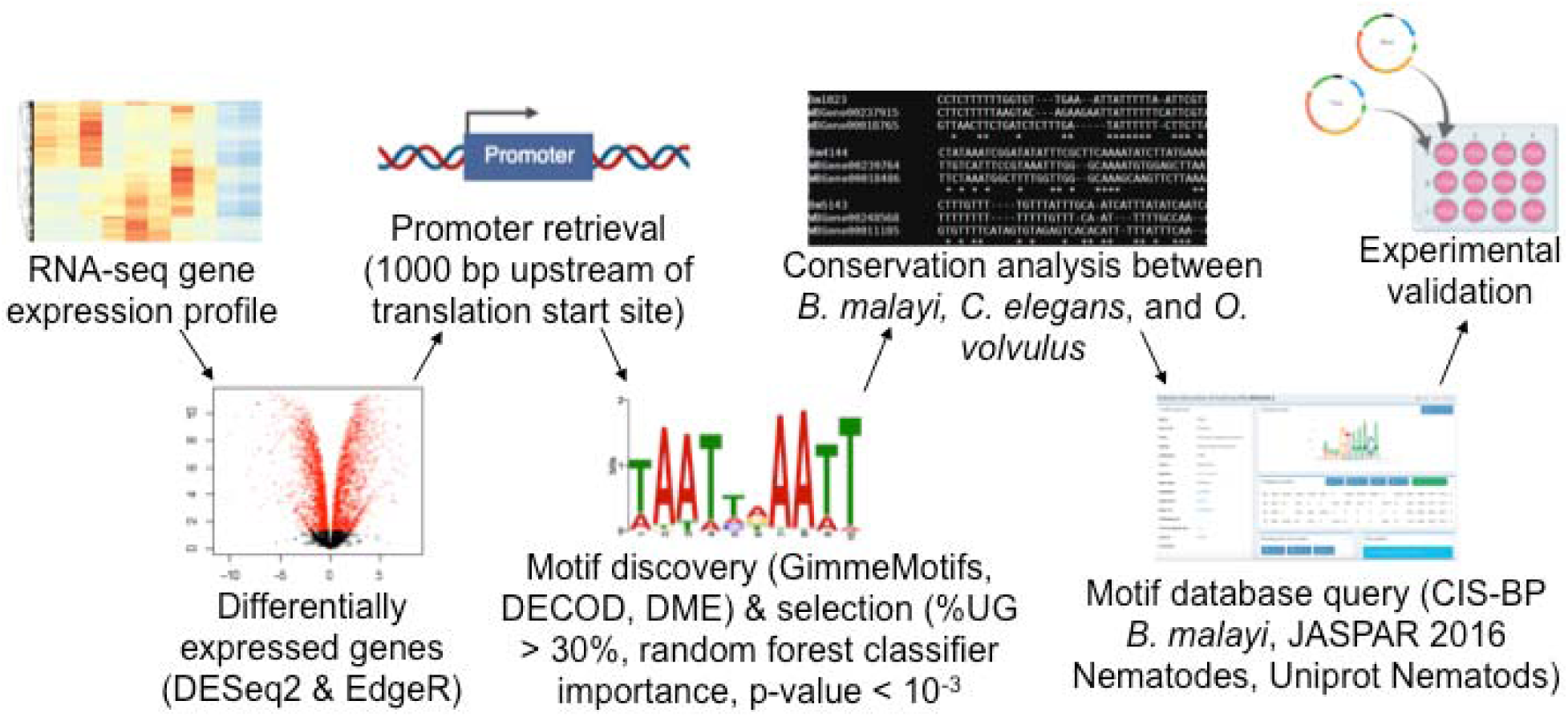
Workflow of promoter motif identification. The six main steps for motif discovery were: 1) generation of an RNA-seq profile, 2) determination of up-regulated genes for every pairwise comparison using DESeq2 (FDR<0.01) and EdgeR (P-value<0.01), 3) promoter retrieval: 1000 bp upstream of the translation start site, 4) ensemble motif discovery using GimmeMotifs (Homer, AMD, BioProspector, MDmodule, MEME, Weeder, GADEM, and Improbizer), DECOD, DME, and selection of enriched motifs: %UG > 30% and random forest classifier feature importance and over-representation p-value < 10^−3^, 5) TFBS conservation analysis between *B. malayi, C. elegans*, and *O. volvulus*. 6) motif database query: CIS-BP *B. malayi*, JASPAR 2016 Nematodes, Uniprot Nematodes. Finally, a subset of those identified motifs were experimentally validated.

To select statistically significant motifs, we first assessed the motifs by a random forest classifier using scikit-learn (Varoquaux et al. 2015). The random forest algorithm uses bootstrap sampling and constructs a decision tree for each sub-sample. To evaluate the motifs, we used both Gini impurity (Gordon et al.1984) and information gain (Quinlan et al. 1993) criteria and retained the union of the resulting top 40 motifs. We further filtered the motifs by foreground coverage (i.e. UG%), removing motifs occurring in less than 30% of the genes up-regulated at that stage of molting. We then used a Z-test to compare the frequency of a motif in the up-regulated genes with the expected frequency in the background promoters. Using a significance level (p-value) cutoff of 10^−3^, we selected 395 motifs.

We retrieved a collection of 163 known nematode transcription factor binding sites (TFBSs) from the MEME suite (http://meme-suite.org/), searching the motif databases JASPAR CORE 2016 nematodes (Mathelier et al. 2016), CIS-BP *Brugia malayi* (Weirauch et al. 2014), and Uniprobe worm (Newburger et al. 2009). We matched the remaining motifs to known TFBSs with TOMTOM (Gupta et al. 2007). If two motifs were matched to the same binding site and they were discovered from the same gene list, we considered them to be redundant and kept the one with the lowest over-representation p-value. This step narrowed our list down to 27 motifs that had matches to known binding sites.

We next performed a conservation analysis amongst nematodes using an adaptation of a published method (Roy et al. 2013). We retrieved orthologous gene information among *B. malayi, C. elegans*, and *O. volvulus* from Wormbase ParaSite Biomart (Howe et al. 2017). We extracted up to 1Kb upstream from the translation start sites for *B. malayi* genes, assuming these regions would contain the promoter. We performed multiple sequence alignments using CLUSTALW2 (Larkin et al. 2007), and defined a motif as conserved if it occurred at the same position in the orthologous promoter region alignment of either *C. elegans* or *O. volvulus*. This step resulted in 12 remaining motifs (**Table 2**) that were (1) enriched (p-value < 10^−3^) and (2) conserved in either *C. elegans* or *O. volvulus*. The frequency of motif occurrence in the putative promoter regions of up-regulated genes ranges from 33% to 94%. The fold enrichment, representing the ratio between motif frequencies in the up-regulated gene promoters and background promoters, ranges from 1.28 to 2.19.

**Table 2:**
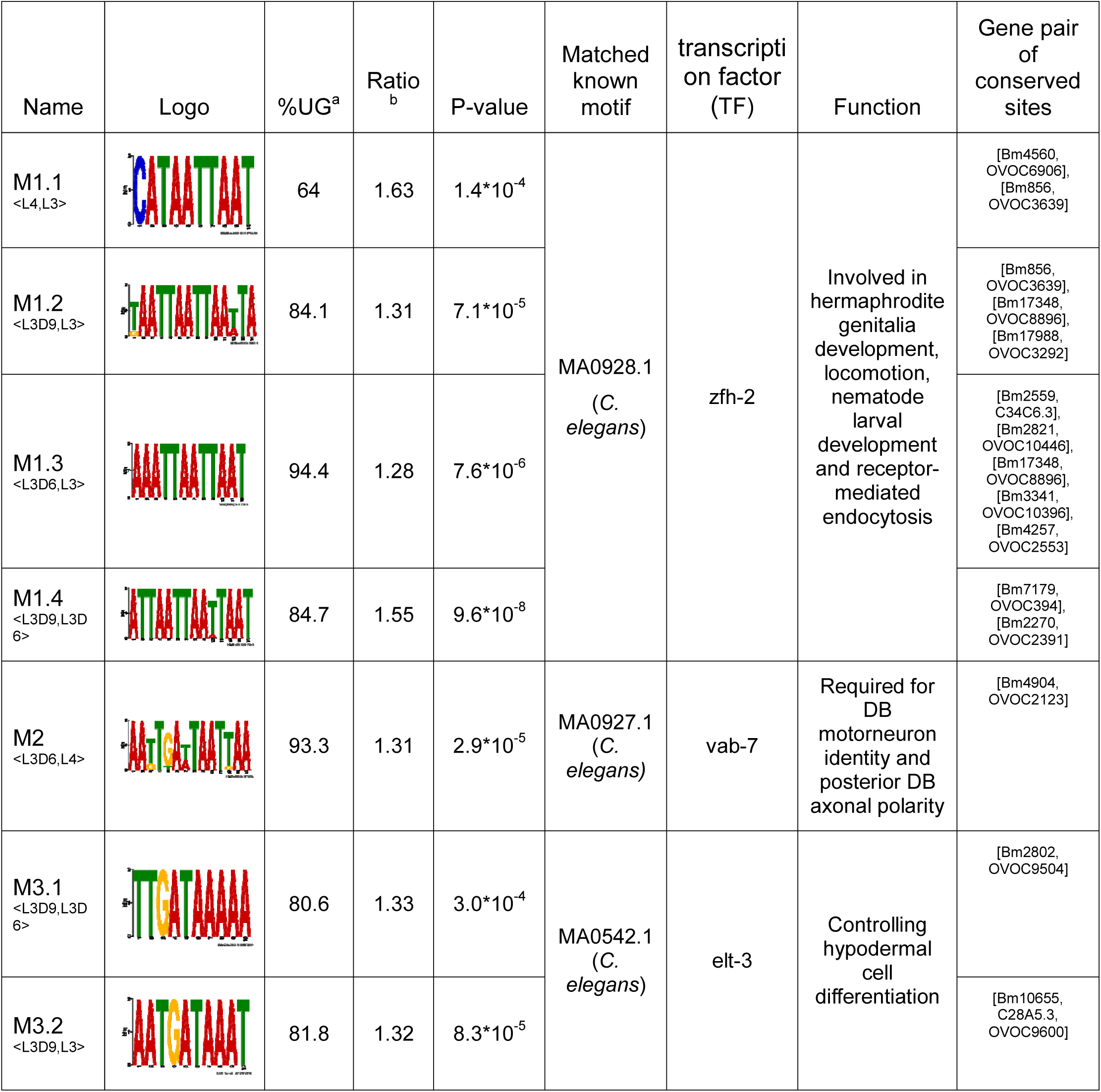
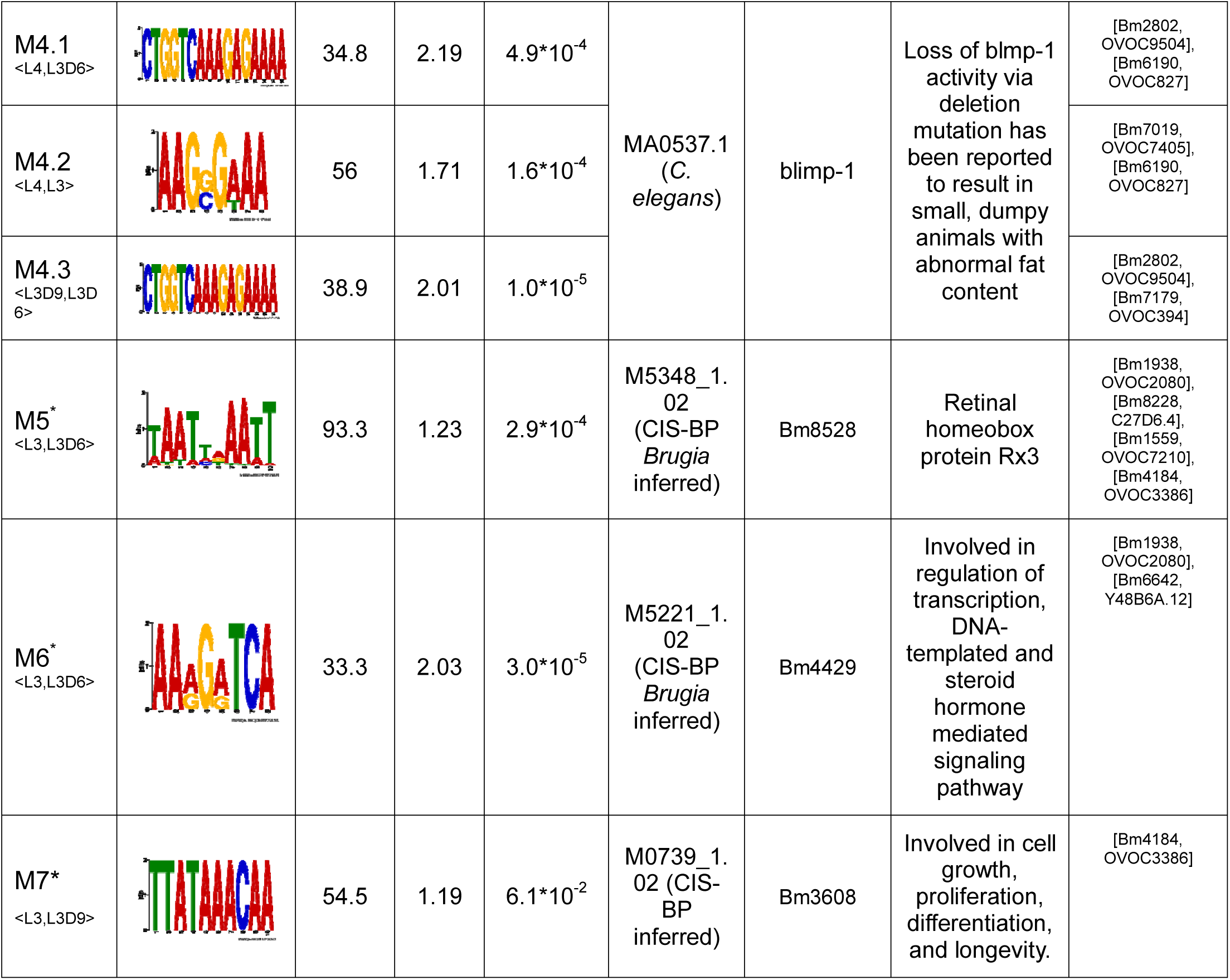
Table of enriched promoter motifs over the L3-L4 molt. Motifs were found to be enriched (p-value < 10^−3^) in the upstream elements of up-regulated genes between different stages of molting and to be conserved (in either *C. elegans* or *O. volvulus*). ^a^Frequency of a motif in up-regulated gene promoters. ^b^Relative frequency of a motif in up-regulated gene promoters vs. background promoters. *These two motifs have been validated experimentally. Note that M7 was included in the experimental validation because it passed 4 out of 5 filters, including foreground coverage filter, random forest filter, known motif filter and conservation filter. However, it was not included in the 12 reported motifs due to its non-significant p-value.

| Name | Logo | %UG<sup>a</sup> | Ratio<sup>b</sup> | P-value | Matched known motif | transcription factor (TF) | Function | Gene pair of conserved sites |
| --- | --- | --- | --- | --- | --- | --- | --- | --- |
| M1.1<br><L4,L3> |  | 64 | 1.63 | 1.4*10<sup>-4</sup> | MA0928.1<br>(*C. elegans*) | zfh-2 | Involved in hermaphrodite genitalia development, locomotion, nematode larval development and receptor-mediated endocytosis | [Bm4560, OVOC6906], [Bm856, OVOC3639] |
| M1.2<br><L3D9,L3> |  | 84.1 | 1.31 | 7.1*10<sup>-5</sup> |  |  |  | [Bm856, OVOC3639], [Bm17348, OVOC8896], [Bm17988, OVOC3292] |
| M1.3<br><L3D6,L3> |  | 94.4 | 1.28 | 7.6*10<sup>-6</sup> |  |  |  | [Bm2559, C34C6.3], [Bm2821, OVOC10446], [Bm17348, OVOC8896], [Bm3341, OVOC10396], [Bm4257, OVOC2553] |
| M1.4<br><L3D9,L3D6> |  | 84.7 | 1.55 | 9.6*10<sup>-8</sup> |  |  |  | [Bm7179, OVOC394], [Bm2270, OVOC2391] |
| M2<br><L3D6,L4> |  | 93.3 | 1.31 | 2.9*10<sup>-5</sup> | MA0927.1<br>(*C. elegans*) | vab-7 | Required for DB motoneuron identity and posterior DB axonal polarity | [Bm4904, OVOC2123] |
| M3.1<br><L3D9,L3D6> |  | 80.6 | 1.33 | 3.0*10<sup>-4</sup> | MA0542.1<br>(*C. elegans*) | elt-3 | Controlling hypodermal cell differentiation | [Bm2802, OVOC9504] |
| M3.2<br><L3D9,L3> |  | 81.8 | 1.32 | 8.3*10<sup>-5</sup> |  |  |  | [Bm10655, C28A5.3, OVOC9600] |
| M4.1<br><L4,L3D6> |  | 34.8 | 2.19 | 4.9*10<sup>-4</sup> | MA0537.1<br>(*C. elegans*) | blimp-1 | Loss of blimp-1 activity via deletion mutation has been reported to result in small, dumpy animals with abnormal fat content | [Bm2802, OVOC9504], [Bm6190, OVOC827] |
| M4.2<br><L4,L3> |  | 56 | 1.71 | 1.6*10<sup>-4</sup> |  |  |  | [Bm7019, OVOC7405], [Bm6190, OVOC827] |
| M4.3<br><L3D9,L3D6> |  | 38.9 | 2.01 | 1.0*10<sup>-5</sup> |  |  |  | [Bm2802, OVOC9504], [Bm7179, OVOC394] |
| M5*<br><L3,L3D6> |  | 93.3 | 1.23 | 2.9*10<sup>-4</sup> | M5348_1.02<br>(CIS-BP *Brugia* inferred) | Bm8528 | Retinal homeobox protein Rx3 | [Bm1938, OVOC2080], [Bm8228, C27D6.4], [Bm1559, OVOC7210], [Bm4184, OVOC3386] |
| M6*<br><L3,L3D6> |  | 33.3 | 2.03 | 3.0*10<sup>-5</sup> | M5221_1.02<br>(CIS-BP *Brugia* inferred) | Bm4429 | Involved in regulation of transcription, DNA-templated and steroid hormone mediated signaling pathway | [Bm1938, OVOC2080], [Bm6642, Y48B6A.12] |
| M7*<br><L3,L3D9> |  | 54.5 | 1.19 | 6.1*10<sup>-2</sup> | M0739_1.02 (CIS-BP inferred) | Bm3608 | Involved in cell growth, proliferation, differentiation, and longevity. | [Bm4184, OVOC3386] |

The 12 selected motifs matched known binding sites for 6 transcription factors in *C. elegans* (**Table 2**), all of which are involved in development, aging, and/ or movement. Motifs M1.1, M1.2, M1.3 and M1.4 matched a zinc-finger protein, zfh-2, which is involved in hermaphrodite genitalia development, locomotion, nematode larval development and receptor-mediated endocytosis. Motif M2 matched vab-7, which is associated with DB motor neuron identity and posterior DB axonal polarity. Motifs M3.1 and M3.2 matched elt-3, which is related to aging (Budovskaya et al. 2008). Motifs M4.1, M4.2, and M4.3 matched blmp-1. M5 matched a homeobox protein, Bm8528, and M6 matched a nuclear receptor Bm4429. The 12 motifs are conserved in either *C. elegans* or *O. volvulus* (**Table 2**, last column). Moreover, the occurrence of M3.2 in the Bm10655 promoter region is conserved in orthologs in both *C. elegans* (promoter of C28A5.3) and *O. volvulus* (promoter of OVOC9600).

The motif analysis reveals how some of the differential expression of different proteases may be orchestrated during the L3 to L4 molt. Motif M1.4 is found in the promoter region of Bm2270 a metalloprotease significantly up-regulated in L3D9 worms. Bm2270 is an ortholog of nas-37 in *C. elegans* and has been shown to be involved in collagen and cuticulin-based cuticle development and ecdysis. Motif M5 is found in the promoter region of Bm1938 and is predicted to encode a serpin. Bm1938 is one of the serpins that was found to be significantly up-regulated in the iL3 larvae.

### L3 stage-specific transcription factor binding motifs can be validated *in vitro*

Three of the motifs (M5, M6, and M7) were chosen for validation based on their enrichment in the promoters of genes up-regulated in the mid to late stages of the L3 to L4 molt. Three separate genes, each containing one chosen motif, were tested. The 1 kbp upstream region of each gene was amplified from *B. malayi* genomic DNA and cloned upstream of the firefly luciferase reporter gene in the expression vector pGL3 Basic (Shu et al. 2003). *B. malayi* L3 were then transfected with the constructs in a co-culture system as previously described (Liu et al. 2018). The parasites were induced to molt in vitro and then assayed for luciferase activity. The number of relative light units (RLUs) observed were normalized to those obtained from parasites transfected in parallel with a construct consisting of the *B. malayi* HSP70 promoter driving the expression of the firefly luciferase reporter (Shu et al. 2018). The experiment was performed with both the□native promoter and a mutant promoter where the nucleotides of the motif had been randomly shuffled. All of the native promoters produced significant amounts of reporter luciferase activity in the molting parasites (ranging from 40%- 70% of the activity produced by the HSP70 construct transfected positive controls; **Fig. 3**). However, when the putative motifs were mutated, the activity in all the promoters tested decreased by 80-90% (**Fig. 3**).

**Figure 3:**
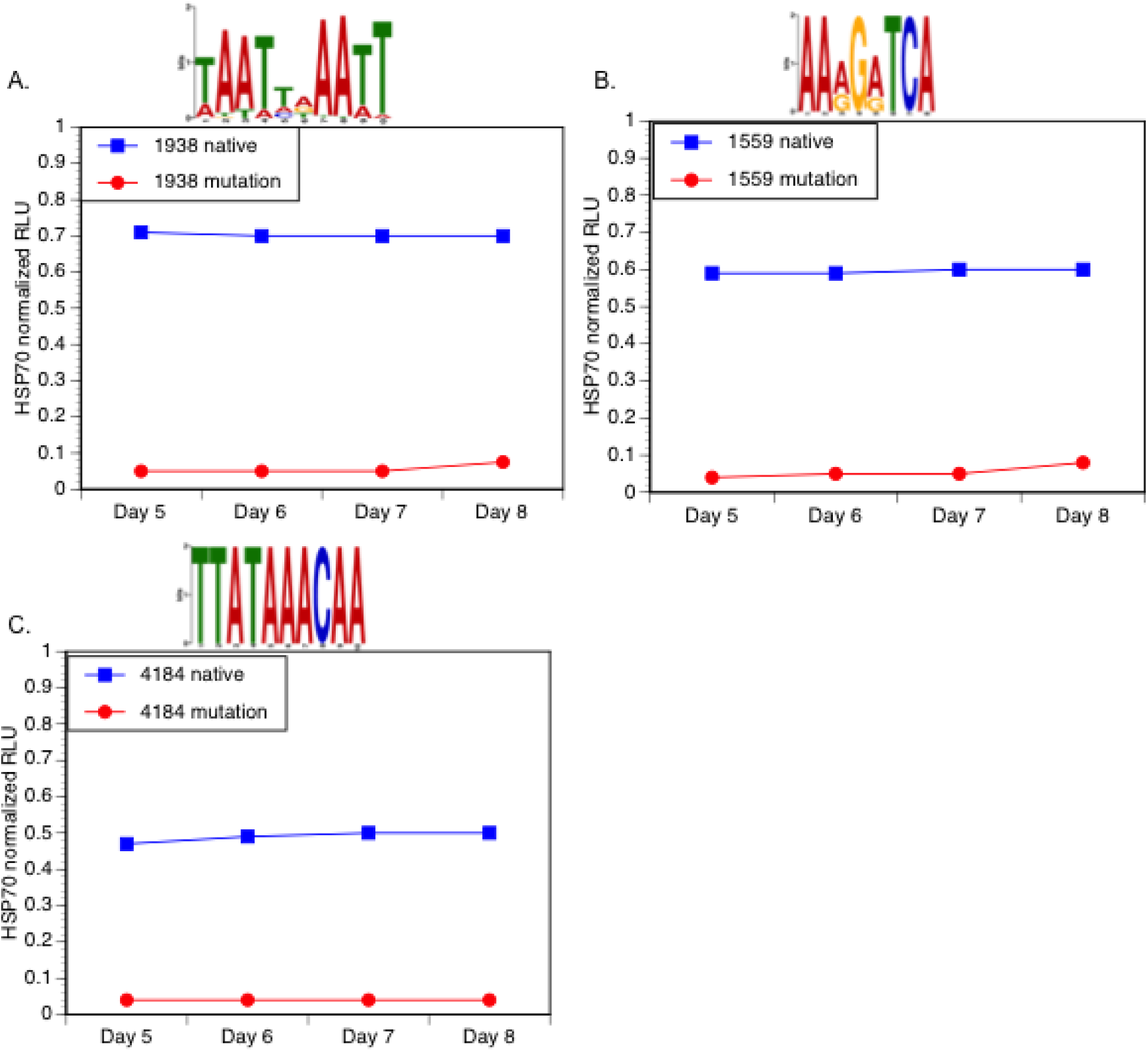
Promoter motif validation. A) Promoter motif validation in L3 worms that were molting in vitro using the native promoter of Bm1559 and a mutated motif M5 version of the same promoter. B) Promoter motif validation using the native promoter of Bm1938 and a mutated motif M6 version of the promoter. C) Promoter motif validation using the native promoter of Bm4184 with a mutated motif M7 version of the promoter. In each panel luciferase activity obtained from the constructs is normalized against parasites transfected with a construct containing the Bm HSP70 promoter driving the expression of the firefly luciferase reporter.

## Discussion

Parasitic nematodes such as *B. malayi* maintain a complicated lifecycle involving both an insect and a mammalian host, and undergo a number of molts. The L3 to L4 molt that occurs immediately upon infection of the mammalian host is of particular interest as it marks the establishment of infection and thus represents an attractive point for drug intervention. Little is known, however, about how *B. malayi* regulates the transitions between these stages. Prior to our study, nothing was known about promoter motifs that regulate developmentally expressed genes in *B. malayi*. Because stage transitions rely on precise transcriptional control through the interaction of transcription factors and their binding sites, we set out to characterize the potential transcription factor binding motifs of these parasites and to identify motifs that contribute to stage-specific expression of genes involved in early worm development in the mammalian host.

We found a number of enriched motifs and were able to define both conserved motifs across molting as well as stage-specific motifs. While some of the motifs we identified are conserved in other nematodes, such as *C. elegans*, a number of motifs represent novel binding sites potentially reflecting the differences in development and the parasitic lifestyle. It is known that molting is regulated by an ecdysone-like response system (Barker et al. 1991; Mhashilkar at al. 2016; Mhashilkar at al. 2016; Warbrick et al. 1993, Lui et al, 2012). Two of the identified motifs and their cognate TFs appear to be related to the ecdysone response. For example, zfh-2, the transcription factor predicted to bind four of our identified motifs, is a common cofactor implicated in ecdysone signaling in *D. melanogaster* (Davis et al. 2011). Blimp-1, the transcription factor predicted to bind three of our identified motifs, is an ecdysone-inducible repressor that is essential for the prepupal development in *Drosophila* (Akagi and Ueda 2011). Validation results suggest that our pipeline is able to identify biologically-relevant motifs involved in molting. This analysis provided biological insight into the development of the parasite as well as the identification of novel drug targets.

Future work needs to be done to expand this analysis across the lifecycle of the nematode at different stages of development, in its different hosts (i.e. human vs. mosquito). While transcriptomic data from these stages exists and can be used to predict motifs, validation at other stages *in vivo* will prove more difficult. However, with recent innovation in filarial transgenics, it is now possible to create stable transgenic parasite lines that will allow functional testing *in vivo* of defined promoter motifs at all life stages of the parasite (Liu et al. 2018).

## Materials and Methods

### Transcriptomic study design

All parasites were obtained from FR3 (Filariasis Research Reagent Resource Center; BEI Resources, Manassas, VA, USA), where they were isolated from infected gerbils (*Meriones unguiculatus*) or mosquitoes (*Aedes aegypti*). Worms were flash-frozen and shipped to the New York Blood Center for processing. For transcriptomic sequencing, infective third-stage larvae (iL3) were recovered from mosquitoes and mammalian stage larvae were recovered from gerbils at 6 and 9 days post infection (dpi). At 6 dpi, larvae are typically undergoing the molt from L3 to L4, while by 9 dpi the molt is complete (Mutafchiev et al. 2014). Data was combined with previously published stages 14 dpi (L4) (Grote et al. 2017).

### RNA isolation, library preparation and sequencing

Total RNA was prepared from *B. malayi* worms as previously described (Grote et al. 2017). RNA was prepared from 3 biological replicates of infective L3 (iL3; 2000 larvae each), 3 replicates of 6 dpi larvae (1500 each) and 2 replicates of 9 dpi larvae (1300 each). *B*. *malayi* worms were homogenized in Trizol (ThermoFisher) using a hand-held pestle in 1.5mL tubes containing the worms. Total RNA was extracted by organic extraction using Trizol and the PureLink RNA mini kit (ThermoFisher) and after being treated with DNaseI (New England Biolabs). Ribosomal RNA (rRNA) depletion was performed using Terminator (Epicentre), a 5’-phosphate-dependent exonuclease that degrades transcripts with a 5’ monophosphate. Libraries were prepared using the NEBNext Ultra II RNA Library Prep Kit for Illumina (New England Biolabs) according to manufacturer instructions. Library quality was assessed using a D1000 ScreenTape Assay (Agilent) prior to sequencing. Library concentrations were assessed using the qPCR library quantification protocol (KAPA biosystems). Libraries were sequenced on the Illumina NextSeq500 platform with 150bp paired-end reads. To minimize the confounding effects of lane-to-lane variation, libraries were multiplexed and sequenced with technical replicates on multiple lanes. Each biological replicate received an average of 135 million mapped reads (PRJNA557263).

### Sequencing alignment and expression analysis

Read quality was assessed using FastQC (Babraham Bioinformatics). Sequence reads from each sample were analyzed with the Tuxedo suite of tools (Kim et al. 2013; Trapnell et al. 2013; Trapnell et al. 2010). Reads were mapped with Tophat2’s Bowtie2-very-sensitive algorithm to the annotated *B. malayi* genome assembly (WormBase.org). The resulting BAM files were then used with HtSeq to obtain raw read counts. Differential gene expression analysis was performed using both DESeq and EdgeR, and the overlapping genes (FDR < 0.01 & P-value < 0.01) were retained. Up-regulated genes were characterized for each pairwise comparison between L3, L3 Day 6 (L3D6), L3 day 9 (L3D9), and L4 worms. For example, <L3,L4> refers to the up-regulated genes in L3 compared to L4. Two pairs of comparisons, <L3D9,L4> and <L4,L3D9>, were dropped due to the limited number of up-regulated genes (<=3), possible due to the L3D9 being actually younger L4. The up-regulated gene lists were filtered using log2 fold change (logFC) with the following thresholds: |logFC| = 7 for <L3D6,L3>, <L3D6,L3D9>, <L3D6,L4>; |logFC| = 4 for <L3D9,L3>, <L3,L3D6>, <L3,L3D9>, <L3,L4>, <L4,L3>; |logFC| = 2.5 for <L3D9,L3D6>, <L4,L3D6>. The reason for varying the threshold was that the number of up-regulated genes in each list varied significantly; for motif discovery tools to search efficiently, the number of sequences were limited to less than one hundred. In total, 10 up-regulated gene lists were used for motif discovery (**Table S1)**.

Potential promoter sequences were retrieved from WormBase ParaSite Biomart (Howe et al. 2016) web interface, capturing the 1000bp upstream of the translation start site for each gene.

### The Emotif Alpha pipeline for regulatory motif identification

The Emotif Alpha pipeline (freely available at: https://github.com/YichaoOU/Emotif_Alpha) was developed to automate motif discovery analysis for the 10 up-regulated gene lists. This pipeline was written in python and was applied to perform all aforementioned motif analyses. The motif discovery step used multiple tools and was run in parallel at the Ohio SuperComputer Center. Motif length search was from 6 to 14. Motif scanning was done using FIMO (Grant et al. 2011) with a default p-value threshold of 10^−4^. We implemented 5 different motif filters. (1) Foreground coverage (i.e., UG%) was defined as the proportion of up-regulated gene promoters containing the given motif. We set a minimal foreground coverage at 30%. (2) Motifs were then filtered by a random forest classifier. The union of the top 40 motifs that resulted from either Gini impurity or information gain criterion was retained. (3) Motif enrichment p-value was calculated using Z-test and the cutoff was 10^−3^. (4) Known motif filter was performed using TOMTOM and a collection of 163 known nematode TFBSs. The motif similarity p-value threshold was 10^−4^. (5) Conservation analysis was performed using a method described in (Roy et al. 2013). Only conserved motifs were kept.

### *In vitro* validation of promoter transcription motifs

The putative TF motifs M5, M6, and M7 were chosen for validation based on their enrichment in the promoters of genes upregulated in the mid to late stages of the L3 to L4 molt. Three different genes, each containing one of the chosen motifs, were used for the validation assay. As previously described in Shu et al. (Shu et al. 2003), we amplified the 1 kbp region upstream of each gene from *B. malayi* genomic DNA and cloned upstream of the firefly luciferase reporter gene in the expression vector pGL3 Basic. We then transfected *B. malayi* L3 larvae with the constructs in a co-culture system as previously described (Liu et al. 2018). The parasites were induced to molt by the addition of ascorbic acid on day 5, and parasites were assayed for luciferase activity on days 5-8, as by day 9 the molting was complete. We normalized the number of RLUs observed to those obtained from parasites transfected in parallel with a construct consisting of the *B. malayi* HSP70 promoter driving the expression of the firefly luciferase reporter (Shu et al. 2018) to control for accumulation of the firefly luciferase over time during the duration of the experiment. We did the experiment with both the□native promoter and a mutant promoter where the nucleotides of the motif had been randomly shuffled (**Table S2**).

## References

Akagi, K., Ueda, H. (2011). Regulatory Mechanisms of Ecdysone-Inducible Blimp-1 Encoding a Transcriptional Repressor that is Important for the Prepupal Development in Drosophila. Development, Growth & Differentiation, 53 (5): 697–703.

Anders, S., & Huber, W. (2010). Differential Expression Analysis for Sequence Count Data. Genome Biology, 11(R106).

Ao, W., Gaudet, J., Kent, W. J., Muttumu, S., & Mango, S. E. (2004). Environmentally Induced Foregut Remodeling by PHA-4/FoxA and DAF-12/NHR. Science, 305(5691), 1743–1746. https://doi.org/10.1126/science.1102216

Bailey, T. L., Williams, N., Misleh, C., & Li, W. W. (2006). MEME: Discovering and Analyzing DNA and Protein Sequence Motifs. Nucleic Acids Research, 34(suppl_2), W369–W373. https://doi.org/10.1093/nar/gkl198

Barker, G. C., J. G. Mercer, H. H. Rees, & R. E. Howells. (1991). The Effect of Ecdysteroids on the Microfilarial Production of Brugia pahangi and the Control of Meiotic Reinitiation in the Oocytes of Dirofilaria immitis. Parasitology Research, 77 (1): 65–71

Bennuru, S., Cotton, J.A., Ribeiro, J.M., Grote, A., Harsha, B., Holroyd, N., Mhashilkar, A., Molina, D.M., Randall, A.Z., Shandling, A.D., Unnasch, T.R., Ghedin, E., Berriman, M., Lustigman, S., Nutman, T.B. (2016). Stage-Specific Transcriptome and Proteome Analyses of the Filarial Parasite Onchocerca volvulus and Its Wolbachia Endosymbiont. mBio, 7(6). PMC5137501.

Breiman, L., Friedman, J., Stone, C., Olshen, R.A. Classification and Regression Trees. Taylor & Francis, Jan 1, 1984.

Budovskaya, Y.V., Wu, K., Southworth, L., Jiang, M., Tedesco, P., Johnson, T., & Kim, S. (2008). An Elt-3/elt-5/elt-6 GATA Transcription Circuit Guides Aging in C. Elegans. Cell, 134 (2): 291–303.

Choi, Y.-J., Ghedin, E., Berriman, M., McQuillan, J., Holroyd, N., Mayhew, G. F., … Michalski, M. L. (2011). A Deep Sequencing Approach to Comparatively Analyze the Transcriptome of Lifecycle Stages of the Filarial Worm, Brugia malayi. PLOS Neglected Tropical Diseases, 5(12), 1–10. https://doi.org/10.1371/journal.pntd.0001409

Conlon, E.M., Liu, X.S., Lieb, J., & Liu, J. (2003). Integrating Regulatory Motif Discovery and Genome-Wide Expression Analysis. Proceedings of the National Academy of Sciences of the United States of America, 100 (6): 3339–44.

Cotton, J. A., Bennuru, S., Grote, A., Harsha, B., Tracey, A., Beech, R., Doyle, S., et al. (2016). The Genome of *Onchocerca volvulus*, Agent of River Blindness. Nature Microbiology, 2 (2). https://doi.org/10.1038/nmicrobiol.2016.216.

Das, M. K., & Dai, H.-K. (2007). A Survey of DNA Motif Finding Algorithms. BMC Bioinformatics, 8(7), S21. https://doi.org/10.1186/1471-2105-8-S7-S21

Davis, M.B., SanGil, I., Berry, G., Olayokun, R., & Neves, L. (2011). Identification of Common and Cell Type Specific LXXLL Motif EcR Cofactors Using a Bioinformatics Refined Candidate RNAi Screen in Drosophila Melanogaster Cell Lines. BMC Developmental Biology, 11 (November): 66.

Desjardins, C. A., et al. (2013). Genomics of Loa loa, a Wolbachia-free Filarial Parasite of Humans. Nature Genetics, 45(5): 495–500.

Dieterich, C., & Sommer, R.J. (2008). A Caenorhabditis Motif Compendium for Studying Transcriptional Gene Regulation. BMC Genomics, 9(1), 30. https://doi.org/10.1186/1471-2164-9-30

Foster, J., Ganatra, M., Kamal, I., Ware, J., Makarova, K., Ivanova, N., Bhattacharyya, A., et al. (2005). The Wolbachia Genome of Brugia Malayi: Endosymbiont Evolution within a Human Pathogenic Nematode. PLoS Biology. 3 (4): e121.

Ghedin, E., Wang, S., Spiro, D., Caler, E., Zhao, Q., Crabtree, J., … Scott, A. L. (2007). Draft Genome of the Filarial Nematode Parasite Brugia malayi. Science, 317(5845), 1756–1760. https://doi.org/10.1126/science.1145406

Gordon, A. D., Breiman, L., Friedman, J.H., Olshen, R.A., & Stone, C.J. (1984). Classification and Regression Trees. Biometrics. https://doi.org/10.2307/2530946.

Gordon, C.A., Jones, M.K., McManus, D.P. (2018) The history of Bancroftian Lymphatic Filariasis in Australasia and Oceania: Is there a Threat of Re-occurance in Mainland Australia? Tropical Medicine and Infectious Disease, 3(2), 58.

Grant C.E., Bailey T.L., Noble W.S. (2011). FIMO: Scanning for Occurrences of a Given Motif. Bioinformatics, 27(7):1017–1018. doi:10.1093/bioinformatics/btr064

Grote, A., Lustigman, S., Ghedin, E. (2017) Lessons from the genomes and transcriptomes of filarial nematodes. Molecular and Biochemical Parasitology, 215:23–29. doi: 10.1016/j.molbiopara.2017.01.004.

Grote, A., Voronin, D., Ding, T., Twaddle, A., Unnasch, T., Lustigman, S., & Ghedin, E. (2017). Defining Brugia Malayi and Wolbachia Symbiosis by Stage-Specific Dual RNA-Seq. PLoS Neglected Tropical Diseases, 11 (3): e0005357.

Guiliano, D.B., Hong, X., McKerrow, J.H., Blaxter, M.L., Oksov, Y., Liu, J., Ghedin, E., Lustigman, S. (2004). A Gene Family of Cathepsin L-like Proteases of Filarial Nematodes are Associated with Larval Molting and Cuticle and Eggshell Remodeling. Mol Biochem Parasitol, 136: 227–42.

Gupta, S., Stamatoyannopoulos, J.A., Bailey, T.L., & Noble, W. (2007). Quantifying Similarity between Motifs. Genome Biology, 8 (2): R24.

Heeringen, S. J. Van, & Veenstra, G. J. C. (2011). GimmeMotifs□: a De Novo Motif Prediction Pipeline for ChIP-sequencing Experiments. Bioinformatics, 27(2), 270–271. https://doi.org/10.1093/bioinformatics/btq636

Heinz, S., Benner, C., Spann, N., Bertolino, E., Lin, Y., Laslo, P., Cheng, J., Murre, C., Singh, H., & Glass, C. (2010). Simple Combinations of Lineage-Determining Transcription Factors Prime Cis-Regulatory Elements Required for Macrophage and B Cell Identities. Molecular Cell. https://doi.org/10.1016/j.molcel.2010.05.004.

Higazi, T. B., DeOliveira, A., Katholi, C. R., Shu, L., Barchue, J., Lisanby, M., & Unnasch, T. R. (2005). Identification of Elements Essential for Transcription in Brugia malayi Promoters. Journal of Molecular Biology, 353(1), 1–13. https://doi.org/https://doi.org/10.1016/j.jmb.2005.08.014

Howe, K. L., Bolt, B. J., Shafie, M., Kersey, P., & Berriman, M. (2017). WormBase ParaSite – a Comprehensive Resource for Helminth Genomics. Molecular and Biochemical Parasitology. https://doi.org/https://doi.org/10.1016/j.molbiopara.2016.11.005

Huggins, P., Zhong, S., Shiff, I., Beckerman, R., Laptenko, O., Prives, C., … Bar-Joseph, Z. (2011). DECOD: Fast and Accurate Discriminative DNA Motif Finding. Bioinformatics, 27(17), 2361–2367. https://doi.org/10.1093/bioinformatics/btr412

Jin, V. X., Apostolos, J., Nagisetty, N. S. V. R., & Farnham, P. J. (2009). W-ChIPMotifs: a Web Application Tool for De Novo Motif Discovery from ChIP-Based High-Throughput Data. Bioinformatics, 25(23), 3191–3193.

Kariuki, M. M., Hearne, L. B., & Beerntsen, B. T. (2010). Differential Transcript Expression between the Microfilariae of the Filarail Nematodes, Brugia malayi and B. pahangi. BMC Genomics 11(1), 225. https://doi.org/10.1186/1471-2164-11-225

Kim, D., Pertea, D.,Trapnell, C., Pimentel, H., Kelley, R., & Salzberg, S. (2013). TopHat2: Accurate Alignment of Transcriptomes in the Presence of Insertions, Deletions and Gene Fusions. Genome Biology, 14 (4): R36.

Larkin, M. A., Blackshields, G., Brown, N. P., Chenna, R., McGettigan, P. A., McWilliam, H., Valentin, F., et al. (2007). Clustal W and Clustal X Version 2.0. Bioinformatics, 23 (21): 2947–48.

Lee, N. K., Azizan, F. L., Wong, Y. S., & Omar, N. (2018). DeepFinder: An Integration of Feature-Based and Deep Learning Approach for DNA Motif Discovery. Biotechnology & Biotechnological Equipment, 1–10.

Li, B.-W., Wang, Z., Rush, A. C., Mitreva, M., & Weil, G. J. (2012). Transcription Profiling Reveals Stage- and Function-Dependent Expression Patterns in the Filarial Nematode Brugia malayi. BMC Genomics, 13(1), 184. https://doi.org/10.1186/1471-2164-13-184

Li, Leping. (2009). GADEM: A Genetic Algorithm Guided Formation of Spaced Dyads Coupled with an EM Algorithm for Motif Discovery. Journal of Computational Biology: A Journal of Computational Molecular Cell Biology, 16 (2): 317–29.

Liu, X., Brutlag, D. L., & Liu, J. S. (2001). BioProspector: Discovering Conserved DNA Motifs in Upstream Regulatory Regions of Co-expressed Genes. Pacific Symposium on Biocomputing, 127–138.

Liu C, Enright T, Tzertzinis G, Unnasch TR. (2012) Identification of Genes Containing Ecdysone Response Elements in the Genome of *Brugia malayi*. Mol Biochem Parasitol, 186:38–43; PMCID: PMC 3501679.

Liu C., Mhashilkar, A.S., Chabanon, J., Xu, S., Lustigman, S., Adams, J.H., et al. (2018) Development of a Toolkit for *piggyBac*-Mediated Integrative Transfection of the Human Filarial Parasite *Brugia malayi*. PLoS Negl Trop Dis, 12(5): e0006509. https://doi.org/10.1371/journal.pntd.0006509

Lustigman S, Zhang J, Liu J, Oksov Y, Hashmi S. (2004). RNA Interference Targeting Cathepsin L and Z-like Cysteine Proteases of Onchocerca volvulus Confirmed their Essential Function during L3 Molting. Mol Biochem Parasitol, 138: 165–70.

Mathelier, A., Fornes, O., Arenillas, D., Chen, C., Denay, G., Lee, J., Shi, W., et al. (2016). JASPAR 2016: A Major Expansion and Update of the Open-Access Database of Transcription Factor Binding Profiles. Nucleic Acids Research, 44 (D1): D110–15.

Mhashilkar, A.S., Adapa, S.R., Jiang, R.H., Williams, S.A., Zaky, W., Slatko, B.E., Luck, A.N., Moorhead, A.R., Unnasch, T.R. (2016) Phenotypic and Molecular Analysis of the Effect of 20-Hydroxyecdysone on the Human Filarial Parasite *Brugia malayi*. Int J Parasitol., 46(5-6):333–41. doi: 10.1016/j.ijpara.2016.01.005.

Mhashilkar, A.S., Vankayala, S.L., Lui, C., Kearns, F., Mehrotra, P., Tzertzinis, G., Palli, S.R., Woodcock, H.L., Unnasch, T.R. (2016) Identification of Ecdysone Hormone Receptor Agonists as a Therapeutic Approach for Treating Filarial Infections. PLoS Neglected Tropical Disease, 10(6):e0004772. doi: 10.1371/journal.pntd.0004772.

Mutafchiev, Y., Bain, O., Williams, Z., McCall, J.W., & Michalski, M.L. (2014). Intraperitoneal Development of the Filarial Nematode Brugia Malayi in the Mongolian Jird (Meriones Unguiculatus). Parasitology Research, 113 (5): 1827–35.

Newburger, D. E., & Bulyk, M. L. (2009). UniPROBE: An Online Database of Protein Binding Microarray Data on Protein-DNA Interactions. Nucleic Acids Research. https://doi.org/10.1093/nar/gkn660.

Pavesi, G., Mereghetti, P., Mauri, G., & Pesole, G. (2004). Weeder Web: Discovery of Transcription Tactor Binding Sites in a Set of Sequences from Co-regulated Genes. Nucleic Acids Research, 32(suppl_2), W199–W203. https://doi.org/10.1093/nar/gkh465

Pedregosa, F., Varoquaux, G., Gramfort, A., Michel, V., Thirion, B., Grisel, O., Blondel, M., Prettenhofer, P., Weiss, R., Dubourg, V., Vanderplas, J., Passos, A., Cournapeau, D., Brucher, M., Perrot, M., Duchesnay, E., Louppe, G. (2012). Scikitlearn: Machine Learning in Python. Journal of Machine Learning Research, 12.

Quang, D., & Xie, X. (2016). DanQ: a Hybrid Convolutional and Recurrent Deep Neural Network for Quantifying the Function of DNA Dequences. Nucleic Acids Research, 44 (11), e107. https://doi.org/10.1093/nar/gkw226

Quinlan, J. R. (1993). Constructing Decision Trees. C4.5. https://doi.org/10.1016/b978-0-08-050058-4.50007-3

Robinson, M. D., McCarthy, D. J., & Smyth, G. K. (2010). edgeR: a Bioconductor Package for Differential Expression Analysis of Digital Gene Expression Data. Bioinformatics, 26. https://doi.org/10.1093/bioinformatics/btp616

Roy, S., Wapinski, I., Pfiffner, J., French, C., Socha, A., Konieczka, J., Habib, N., Kellis, Thompson M., & Regev, A. (2013). Arboretum: Reconstruction and Analysis of the Evolutionary History of Condition-Specific Transcriptional Modules. Genome Research, 23 (6): 1039–50.

Shi, J., Yang, W., Chen, M., Du, Y., Zhang, J., & Wang, K. (2011). AMD, an Automated Motif Discovery Tool Using Stepwise Refinement of Gapped Consensuses. PloS One, 6 (9): e24576.

Shu, L., Katholi, C.R., Higazi, T., & Unnasch, T.R. (2003). Analysis of the Brugia Malayi HSP70 Promoter Using a Homologous Transient Transfection System. Molecular and Biochemical Parasitology, 128 (1): 67–75.

Smith, A. D., Sumazin, P., & Zhang, M. Q. (2005). Identifying tissue-selective transcription factor binding sites in vertebrate promoters. Proceedings of the National Academy of Sciences of the United States of America. https://doi.org/10.1073/pnas.0406123102

Thijs, G., Marchal, K., Lescot, M., Rombauts, S., Moor, B. D., Rouze, P., & Moreau, Y. (2002). A Gibbs Sampling Method to Detect Overrepresented Motifs in the Upstream Regions of Co-expressed Genes. Journal of Computational Biology, 9. https://doi.org/10.1089/10665270252935566

Trapnell, C., Hendrickson, D.G., Sauvageau, M., Goff, L., Rinn, J.L., & Pachter, L. (2013). Differential Analysis of Gene Regulation at Transcript Resolution with RNA-Seq. Nature Biotechnology, 31 (1): 46–53.

Trapnell, C., Williams, B.A., Pertea, G., Mortazavi, A., Kwan, G., van Baren, M.J., Salzberg, S.L., Wold, B.J., & Pachter, L. (2010). Transcript Assembly and Quantification by RNA-Seq Reveals Unannotated Transcripts and Isoform Switching during Cell Differentiation. Nature Biotechnology, 28 (5): 511–15.

Varoquaux, G., Buitinck, L., Louppe, G., Grisel, L., Pedregosa, F., & Mueller, A. (2015). Scikit-Learn. GetMobile: Mobile Computing and Communications. https://doi.org/10.1145/2786984.2786995.

Warbrick, E.V., Barker, G.C., Rees, H.H., Howells, R.E. (1993). The Effect of Invertebrate Hormones and Potential Hormone Inhibitors on the Third Larval Moult of the Filarial Nematode, *Dirofilaria immitis*, in vitro. Parasitology, 107: 459–463.

Weirauch, M.T., Yang, A., Albu, M., Cote, A.G., Montenegro-Montero, A., Drewe, P., Najafabadi, H.S., et al. (2014). Determination and Inference of Eukaryotic Transcription Factor Sequence Specificity. Cell, 158 (6): 1431–43.

Xu, S., Lui, C., Tzertzinis, G., Ghedin E., Evans, C., Kaplan, R., Unnasch T. (2011). *In Vivo* Transfection of Developmentally Competent *Brugia malayi* Infective Larvae. International Journal for Parasitology, 41(3-4): 355–362.

Zang X., Maizels R.M. (2001). Serine Proteinase Inhibitors from Nematodes and the Arms Race between Host and Pathogen. Trends Biochemical Science, 26: 191–197.

